# Transcriptomic and proteomic analysis of marine nematode *Litoditis marina* acclimated to different salinities

**DOI:** 10.1101/2021.11.16.468782

**Authors:** Yusu Xie, Liusuo Zhang

**Affiliations:** CAS and Shandong Province Key Laboratory of Experimental Marine Biology, Institute of Oceanology, Chinese Academy of Sciences, Qingdao, 266071, China; Laboratory of Marine Biology and Biotechnology, Qingdao National Laboratory for Marine Science and Technology, Qingdao, 266237, China; Center for Ocean Mega-Science, Chinese Academy of Sciences, 7 Nanhai Road, Qingdao, 266071, China

**Keywords:** salinity, euryhalinity, hyposaline, hypersaline, marine nematode, *Litoditis marina*

## Abstract

Salinity is a critical abiotic factor for all living organisms. The ability to adapt to different salinity environments determines an organism’s survival and ecological niches. *Litoditis marina* is a euryhaline marine nematode widely distributed in coastal ecosystems all over the world, although numerous genes involved in its salinity response have been reported, the adaptive mechanisms underlying its euryhalinity remain unexplored. Here, we utilized worms which have been acclimated to either low salinity or high salinity conditions and evaluated their basal gene expression at both transcriptomic and proteomic levels. We found that several conserved regulators, including osmolytes biosynthesis genes, transthyretin-like family genes, V-type H^+^-transporting ATPase and potassium channel genes, were involved in both short-term salinity stress response and long-term acclimation processes. In addition, we identified genes related to cell volume regulation, such as actin regulatory genes, Rho family small GTPases and diverse ion transporters, might contribute to hyposaline acclimation, while the glycerol biosynthesis genes *gpdh-1* and *gpdh-2* accompanied hypersaline acclimation in *L. marina*. Furthermore, *gpdh-2* might play an essential role in transgenerational inheritance of osmotic stress protection in *L. marina* as in its relative nematode *Caenorhabditis elegans*. Hereby, this study paves the way for further in-depth exploration on adaptive mechanisms underlying euryhalinity, and may also contribute to the studies of healthy ecosystems in the context of global climate change.

## Introduction

Salinity is an important abiotic environmental factor, which affects survival, growth, reproduction and ecological distribution of living organisms. Efficient sensation, response and adaption to their external salinity environment is vital for all living individuals. The imbalance of salt intake also affects human health, which is associated with a variety of cardiovascular diseases and other physiological pathologies (Caudarella et al., 2009; Rust and Ekmekcioglu, 2017; He and MacGregor, 2018). Therefore, studies on osmoregulation mechanisms have always been the hot topic for life.

In the process of response and adaptation to salinity changes in the surrounding environment, certain universal strategies are applied by different organisms, yet with diversities in detailed regulation. Studies using the brewer’s yeast as a model, have demonstrated that osmotic sensation and transduction within a single eukaryotic cell can be highly complex, acting in parallel pathways, and often cross-communicate with other signaling processes (Brewster and Gustin, 2014; Hohmann, 2015). Yeast cells accumulate glycerol as compatible osmolyte under hyper-osmotic stress, and the high osmolarity glycerol (HOG) response pathway controls glycerol accumulation at various levels including the activation of mitogen-activated protein kinase (MAPK) pathway genes (Saito and Posas, 2012; Hohmann, 2015; Tatebayashi et al., 2020). Upon acute osmotic shocks, cell volume changes rapidly. Hyperosmotic shock causes cell shrinkage, while hypoosmotic stress leads to cell swelling. Along with volume perturbation, biophysical properties of cell membrane, physiological state in the cytosol, and DNA structure as well as gene expression in the nucleus will be affected (Hoffmann et al., 2009; Saito and Posas, 2012). It is extensively known that cytoskeleton such as actin (Rizoli et al., 2000; Carton et al., 2003; Di Ciano-Oliveira et al., 2006; Tamma et al., 2007), signaling pathways such as Rho family small GTP binding proteins (Raitt et al., 2000; Carton et al., 2003; Di Ciano-Oliveira et al., 2003) and MAPK (Posas and Saito, 1997; Uhlik et al., 2003; Tanaka et al., 2014), membrane transporters such as water channel aquaporins (Liu et al., 2006; Soveral et al., 2006; Galizia et al., 2008; Furukawa et al., 2009) and a variety of sodium, chloride and potassium related ion channels (Lauf and Theg, 1980; Liedtke et al., 2000; Garnovskaya et al., 2003; Hoffmann et al., 2009), together with organic osmolytes such as glycerol, *myo*-inositol, taurine, and methylamines (Yancey et al., 1982; Yancey, 2005; Burg and Ferraris, 2008), play critical roles in the process of osmotic regulation. These components are also involved in ensuing adaptive regulation of cells under long-term osmotic stresses (Hoffmann et al., 2009; Saito and Posas, 2012). In multicellular organisms with more complicated structures, osmoregulation is conducted mainly by osmoregulatory tissues and organs, for instance, gill in crustacean and fish, kidney in mammals, which are even regulated by their neuroendocrine systems (Sinke and Deen, 2011; Kültz, 2012; Thabet et al., 2017; Niu et al., 2020; Pool et al., 2020; Vij et al., 2020). Moreover, disruption of important members in osmoregulatory process are reported contributing to diverse human diseases (Wilson et al., 2001; Sinke and Deen, 2011; Meor Azlan and Zhang, 2020; Silverman et al., 2020; Li et al., 2021). However, the precise mechanisms of osmotic sensation, signal transduction and adaptation remain poorly defined in invertebrate animals.

Marine nematode *L. marina* has emerged as an excellent invertebrate system for osmoregulation studies, it is a euryhaline nematode, inhabiting widely in the littoral zone of coasts and estuaries all over the world (Xie et al., 2020; Xie et al., 2021; Zhao et al., 2021). In nematodes, the hypodermis and its outer cuticle, the excretory system which is composed of the “H”-shaped excretory cell, the duct cell and the pore cell, as well as the intestine are important osmoregulatory tissues, as well documented by studies in *C. elegans* (Choe and Strange, 2007b; Choe, 2013). Extensive studies in *C. elegans* as well as some extremophilic nematodes have revealed a set of important osmoregulatory genes in nematodes, for instance, metabolic genes related to osmolytes glycerol and trehalose synthesis and accumulation (Lamitina et al., 2006; Adhikari et al., 2009; Burton et al., 2017; Urso and Lamitina, 2021), ion transport related genes like transient receptor potential cation channel TRP family genes and chloride channel genes (Choe, 2013), aquaporin water channel genes (Huang et al., 2007; Igual Gil et al., 2017), extracellular proteins including some collagens and *osm* genes (Lamitina et al., 2006; Wheeler and Thomas, 2006; Rohlfing et al., 2011; Choe, 2013; Dresen et al., 2015; Dodd et al., 2018; Urso and Lamitina, 2021), as well as genes related to MAPK, WNK-1/GCK-3, Notch and insulin-like signaling pathways (Lamitina and Strange, 2005; Choe and Strange, 2007a; Choe, 2013; Burton et al., 2017; Burton et al., 2018; Dodd et al., 2018; Urso and Lamitina, 2021). However, osmoregulatory studies in the terrestrial nematode *C. elegans* were carried out mostly under hyperosmotic conditions. With a simple body structure, a short lifecycle, an annotated reference genome and applicable gene editing manipulation in *L. marina*, this euryhaline marine nematode can thus be used as an attractive experimental system in studying regulatory and adaptive mechanisms underlying both hyperosmotic and hypoosmotic conditions to delineate the molecular mechanisms underlying euryhalinity.

Previously, we applied *L. marina* and challenged them by either low or high salinity stresses, identified a broad range of salinity responding genes at the transcriptional level (Xie et al., 2021). We demonstrated that transthyretin-like family genes and heat shock protein genes were presumably general regulators involved in *L. marina*’s damage control mechanism in response to different salinity stresses. Particularly, unsaturated fatty acids biosynthesis related genes and certain cytoskeleton related genes probably play an important role in response to hyposaline stress, whereas glycerol biosynthesis gene and cuticle related collagen genes were involved in hypersaline stress response (Xie et al., 2021).

To further explore the adaptive mechanisms underlying its euryhalinity, *L. marina* worms were acclimated to either lower salinity or higher salinity culture conditions in the present study, and then transcriptomic and proteomic analysis were performed to study the basal mRNA and protein differences among worms acclimated to different salinity conditions. We found that there are conserved regulators, such as osmolytes biosynthesis genes, various cellular stress response genes including transthyretin-like family genes, as well as cell volume regulation related genes like V-type H^+^-transporting ATPase and potassium channel gene, involved in both short-term salinity stress response and long-term acclimation processes. In addition, we demonstrated that trehalose biosynthesis genes and many genes related to cell volume regulation, such as actin regulatory genes, Rho family small GTPases and diverse ion transporters, presumably contribute to hyposaline acclimation, while genes related to glycerol biosynthesis most probably accompanies hypersaline acclimation in *L. marina*. Furthermore, our results also support the idea of a specific role of the glycerol biosynthesis gene *gpdh-2* in transgenerational inheritance of osmotic stress protection in worms. Many identified genes have orthologues in human which were implicated in various diseases. Nowadays, gradual climate change has already accelerated rises in sea level, which will lead to decreasing of ocean salinity while increased salinization of coastal areas, as a result, more species will encounter either hyposaline or hypersaline stresses. Hereby, our current data will not only pave the way for further in-depth exploration on adaptive mechanisms underlying euryhalinity, but also will provide insights into protection and administration of ecosystems which are stressed by gradual climate change.

## Materials and Methods Worms

Worms acclimated to seawater salinity environment (S30 group): the wild strain of marine nematode *L. marina*, HQ1, was cultured on <u>s</u>ea<u>w</u>ater-NGM (SW-NGM) agar plates prepared with natural seawater (**Supplementary Table 1**), as described previously (Xie et al., 2020; Xie et al., 2021). The salinity of seawater used for this condition is 3%. *E. coli* OP50 was applied as a food source. Worms cultured under this condition were propagated for about 3.5 years till this study.

Worms acclimated to low salinity environment (S3 group): HQ1 worms were transferred from SW-NGM agar plates to normal NGM agar plates containing 0.3% NaCl (**Supplementary Table 1**), which is the standard culture condition for the terrestrial nematode *C. elegans*. OP50 was applied as a food source. Worms cultured under this condition were propagated for about 1.5 years till this study.

Worms acclimated to a higher salinity environment (S50 group): HQ1 worms were transferred from SW-NGM agar plates to the <u>a</u>rtificial <u>s</u>ea<u>w</u>ater-NGM (ASW-NGM) agar plates containing 5% sea salt (**Supplementary Table 1**). ASW-NGM agar plates were prepared by artificial Sea Salt (Instant Ocean). OP50 was applied as the food source. Worms cultured under this condition were propagated for about 3 months (more than 10 generations) till this study.

All worms were maintained at 20°C in the laboratory.

### Developmental Analysis under Different Salinity Conditions

For developmental analysis, 100 newly hatched *L. marina* L1 larvae were transferred onto each indicated 35 mm dimeter agar plates seeded with 30 μl OP50. The number of adult worms was scored every 24 h.

Data represent the average of three replicates. Statistical significance was determined using the two-tailed Student’s *t*-test between two groups. *P* value < 0.05 was considered statistically significant.

For egg laying time analysis, the earliest observed egg laying time on each assay plate was recorded, 100 L1 larvae per plate in six replicates for each condition.

### Lifespan Assay

Worms were cultured under optimal growth conditions for at least 3 generations before lifespan assay. The lifespan assay was performed starting at day 1 of adulthood as described previously (Lee et al., 2010; Xie et al., 2020), with minor modification. In brief, 35 mm diameter assay plates seeded with 30 μl OP50 were prepared every day. Forty L4 females were transferred to each assay plate, incubated at 20 °C. The number of live and dead worms was determined using a dissecting microscope every 24 h. Alive worms were transferred to fresh OP50-seeded agar plates daily. Three replicates were analyzed. Worms were scored as dead if no response was detected after prodding with a platinum wire. Dead worms on the wall of the plate were not counted.

Statistical analysis of the average lifespan for worms acclimated to each salinity condition was performed. Data represent the average of three replicates. The comparisons between two groups were performed using the two-tailed Student’s *t*-test. *P* value < 0.05 was considered statistically significant.

### Synchronized L1 Larvae Collection for Each Salinity Condition

The synchronized L1 larvae under each salinity condition were collected as previously reported by Xie et al. (Xie et al., 2021). Worms, acclimated to S3, S30 and S50 conditions, were allowed to lay eggs overnight at 20°C. Eggs were washed off and collected using corresponding suitable solution, specifically, M9 buffer for S3 group, filtered sterile seawater for S30 group, and filtered sterile artificial seawater containing 5% Sea Salt (Instant Ocean) for S50 group. Then, the eggs were treated with Worm Bleaching Solution (Sodium hypochlorite solution : 10 M NaOH : H_2_O = 4 : 1 : 10, prepared in terms of volume ratio) at room temperature for 1.5 min. Eggs were then washed twice with corresponding suitable solution. The above clean eggs hatched overnight and developed into L1 larvae in corresponding suitable solution at 20°C. The synchronized L1 larvae were collected by filtration using 500 grid nylon filter with 25 μm mesh size. The samples were frozen immediately in liquid nitrogen.

### RNA-seq Analysis

Total RNA was extracted using Trizol (Invitrogen). With three biological replicates for each group, a total of nine RNA libraries were prepared with 3 μg RNA using NEBNext® UltraTM RNA Library Prep Kit for Illumina® (NEB, USA) following manufacturer’s recommendations. Then, RNA libraries were sequenced on an Illumina NovaSeq 6000 platform and 150 bp paired-end reads were generated.

Firstly, Clean data were obtained by removing reads containing sequencing adaptors, reads having poly-N and low-quality ones from raw data. The minimum of base score Q20 was over 97.79% and Q30 was over 93.64%. Then, the clean data were aligned to the *L. marina* reference genome (Xie et al., 2020) by Hisat2 (v2.0.5, with the default parameters) (Kim et al., 2015), with mapping ratio from 68.57% to 70.70% (**Supplementary Table 2**). New transcripts for novel genes were predicted and assembled by StringTie (v1.3.3b, with the default parameters) (Pertea et al., 2015), then annotated with Pfam, SUPERFAMILY, Gene Ontology (GO) and KEGG databases. Briefly, the functional annotation was performed using InterProScan v5.17-56.0 (Jones et al., 2014) by searching against publicly available databases Pfam (http://pfam.xfam.org/), SUPERFAMILY (http://supfam.org), and GO (http://www.geneontology.org/), with an E value cutoff of 1e-5. KEGG function (Kanehisa and Goto, 2000) was assigned using KOBAS 3.0 (Xie et al., 2011) by best hit (with an E value cutoff of 1e-5) to KEGG database (http://www.genome.jp/kegg/). Further, the reads numbers mapped to each gene were analyzed using featureCounts (v1.5.0-p3, with parameter -Q 10 -B -C) (Liao et al., 2014), and FPKM (expected number of Fragments Per Kilobase of transcript sequence per Millions base pairs sequenced of each gene) was calculated based on the length of the gene and reads count mapped to this gene, which was used for estimating gene expression levels. Hierarchical clustering for 9 samples was performed using the pheatmap package in R and can be visualized in Supplementary Figure 1, indicated the reliable sample preparation. Subsequently, differential expression analysis of two conditions was performed using the DESeq2 R package (v1.16.1) (Love et al., 2014). The resulting *P* values were adjusted using the Benjamini and Hochberg’s approach for controlling the false discovery rate. Genes with an adjusted *P* value (padj) < 0.05 found by DESeq2 were assigned as differentially expressed. Moreover, GO enrichment analysis and KEGG pathway enrichment analysis for differentially expressed genes (DEGs) were achieved by clusterProfiler R package (v3.4.4), an adjusted *P* value (padj) < 0.05 was considered significantly enriched. GeneRatio was defined as the ratio of the number of differential genes annotated to the GO term or on the KEGG pathway to the total number of differential genes, respectively.

### Proteomic Analysis

Worm sample was sonicated three times on ice using a high intensity ultrasonic processor (Scientz) in lysis buffer (8 M urea, 1% protease inhibitor cocktail), with three biological replicates for each group. The remaining debris was removed by centrifugation at 12,000 g at 4 °C for 10 min. Then, the supernatant was collected and the protein concentration was determined with a BCA kit (Beyotime Biotechnology) according to the manufacturer’s instructions. Protein samples were digested with trypsin (Promega) at 1:50 trypsin-to-protein mass ratio overnight. After being desalted by Strata X C18 SPE column (Phenomenex) and vacuum-dried, peptides were labelled with a tandem mass tags (TMT) kit (ThermoFisher Scientific) according to the manufacturer’s protocol.

Next, the TMT labeled peptides were fractionated by high-pH reverse-phase HPLC using Agilent 300 Extend C18 column (5 μm particles, 4.6 mm ID, 250 mm length). Briefly, peptides were first separated with a gradient of 8% to 32% acetonitrile (pH 9.0) over 60 min into 60 fractions. Then, the peptides were combined into 9 fractions and dried by vacuum centrifugation.

Further, peptides were identified and quantified by liquid chromatography-tandem mass spectrometry (LC-MS/MS). Briefly, the tryptic peptides were dissolved in solvent A (0.1% formic acid in 2% acetonitrile), directly loaded onto a home-made reversed-phase analytical column (75 μm ID, 15 cm length). The gradient was comprised of an increase from 9% to 22% solvent B (0.1% formic acid in 90% acetonitrile) over 40 min, 22% to 32% in 14 min and climbing to 80% in 3 min then holding at 80% for the last 3 min, all at a constant flow rate of 450 nL/min on an EASY-nLC 1200 UPLC system (ThermoFisher Scientific). The peptides were subjected to nanospray ionization (NSI) source followed by tandem mass spectrometry (MS/MS) in Q Exactive HF-X (ThermoFisher Scientific) coupled online to the UPLC. The electrospray voltage was set as 2.2 kV. The m/z scan range was 350 to 1400 for full scan, and intact peptides were detected in the Orbitrap at a resolution of 120,000. Peptides were then selected for MS/MS using normalized collisional energy (NCE) setting as 28 and the ion fragments were detected in the Orbitrap at a resolution of 30,000. A data-dependent procedure that alternated between one MS scan followed by 20 MS/MS scans with 15 s dynamic exclusion. Automatic gain control (AGC) was set at 1E5. The fixed first mass was set as 100 m/z.

The resulting MS/MS data were processed using Maxquant search engine (v1.5.2.8). Tandem mass spectra were searched against the 17661 protein database of *L. marina* (Xie et al., 2020) concatenated with reverse decoy database. Trypsin/P was specified as cleavage enzyme allowing up to 2 missing cleavages, five modifications per peptide. The mass tolerance for precursor ions was set as 10 ppm in the first search and 5 ppm in the main search, and the mass tolerance for fragment ions was set as 0.02 Da. Carbamidomethyl on Cys was specified as fixed modification, and acetylation on protein N-terminal, oxidation on Met, deamidation on Asn and Gln were specified as variable modifications. Minimum peptide length was set at 7. The quantitative method was set to TMT 10plex, and the false discovery rate (FDR) for protein identification was adjusted to < 1%. All the other parameters in MaxQuant were set to default values. A *P* value < 0.05 from t-tests and a fold change > 1.3 or < 1/1.3 were set as the thresholds for significantly differentially expressed proteins (DEPs).

A total of 306,324 spectrums were acquired, of which 78,015 unique spectrums were obtained, and a total of 45,669 peptides were identified (**Supplementary Table 3**). The length distribution analysis of peptides showed that most of them consisted of 7-20 amino acids, which is in accordance with the quality control requirements (**Supplementary Figure 2**). Subsequently, hierarchical clustering for 9 samples were shown in Supplementary Figure 3, indicated the reliable sample preparation.

Moreover, different databases were selected for protein functional annotation. GO annotation proteome was derived from the UniProt-GOA database (http://www.ebi.ac.uk/GOA/). If the proteins were not annotated by UniProt-GOA database, the InterProScan (https://www.ebi.ac.uk/interpro) would be applied to annotate by protein sequence alignment method, which was also used for protein domain functional description. KEGG online service tools KAAS was utilized to predict the pathways in which the identified proteins were involved. Then the annotation results were mapped on the KEGG pathway database using KEGG mapper. Subcellular localizations of the protein were predicted by wolfpsort (http://wolfpsort.seq.cbrc.jp/). COG annotation of the protein was achieved by eggnog-mapper software (v2.0) with the default parameters.

Additionally, enrichment analysis of GO and KEGG pathway was conducted for DEPs by Python using an in-house script, two-tailed Fisher’s exact test was employed to test the enrichment of DEPs against the background of all identified proteins, a corrected *P* value < 0.05 was considered significantly enriched. Ratio was defined as the ratio of the number of differential proteins annotated to the GO term or on the KEGG pathway to the total number of differential proteins, respectively.

## Results

### *L. marina* is a Typical Euryhaline Marine Nematode

In the laboratory, marine nematode *L. marina* HQ1 was normally cultured on SW-NGM agar plates prepared by seawater with a salinity of 3%, and this condition was referred to as “S30” in this paper. Basic developmental characteristics were tested in the laboratory, including the adulthood percentage, the earliest egg laying time, as well as the lifespan. Under S30 condition, although only 3% newly hatched L1 larvae could develop into adult stage within 3 days at 20°C, the adulthood percentage was as high as 92% within a week (**Figure 1A**). The earliest observed egg laying time was around 84 h post L1 stage (**Figure 1B**). Moreover, L4 worms could live as long as 29 days (**Figure 1C**), with an average lifespan of about 16 days (**Figure 1D**).

**FIGURE 1.**
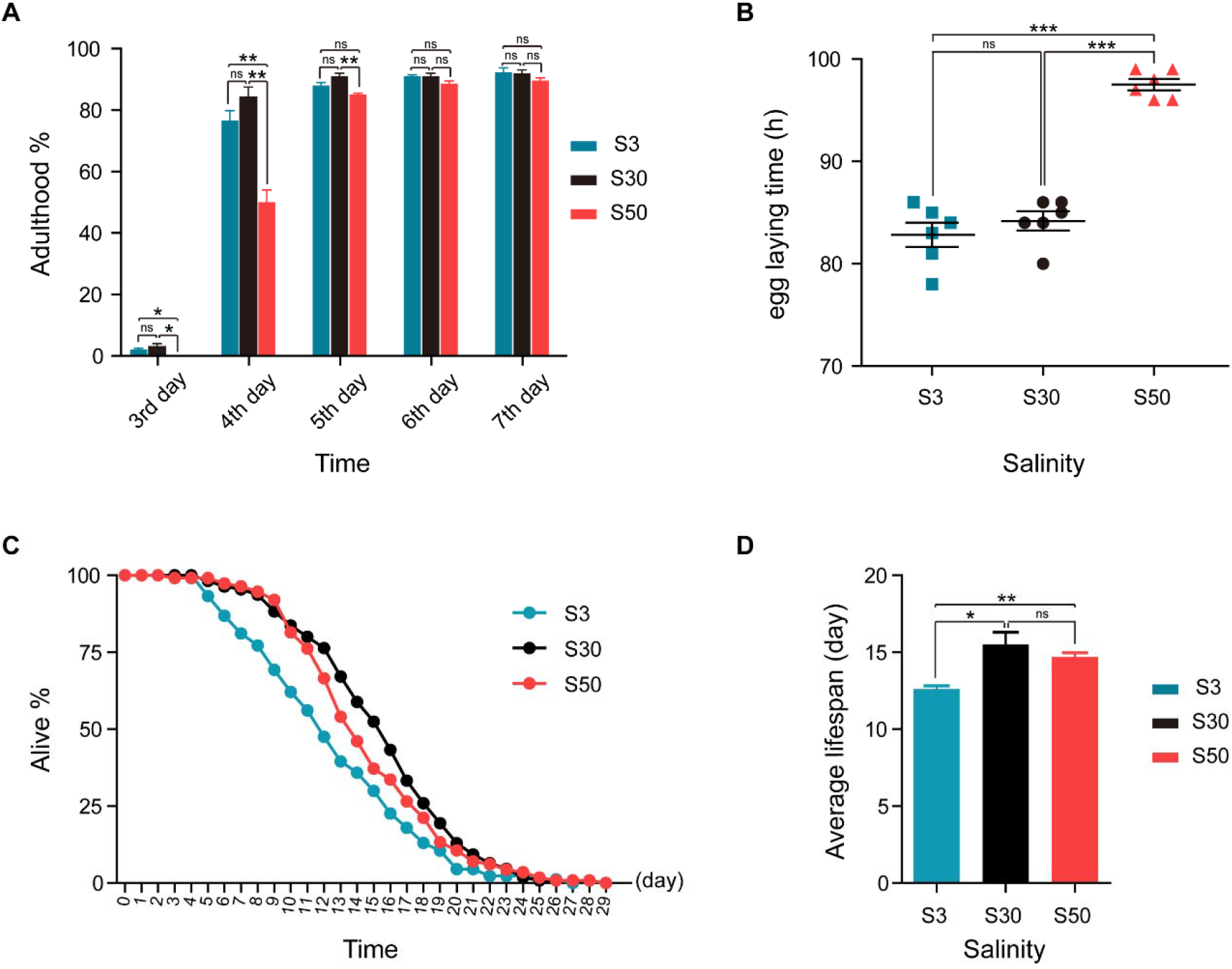
Developmental characterization and lifespan of *L. marina* acclimated to different salinity conditions. (**A**) For developmental analysis, 100 newly hatched L1s were transferred onto each indicated 35 mm dimeter agar plates seeded with 30 μl OP50. The number of adult worms was scored every day. (**B**) Egg laying time of *L. marina* acclimated to different salinity conditions. One hundred newly hatched L1s were transferred onto each indicated 35 mm dimeter agar plates seeded with 30 μl OP50. The earliest observed egg laying time on each plate was recorded. Six replicates were performed for each experimental condition. (**C**) For lifespan assay, 40 L4 females were transferred to each assay plate, incubated at 20°C. The number of live and dead worms was determined using a dissecting microscope every 24 h. Alive worms were transferred to fresh OP50-seeded plates daily. Three replicates were analyzed. Worms were scored as dead if no response was detected after prodding with a platinum wire. Dead worms on the wall of the plate were not counted. (**D**) Average lifespan of *L. marina* acclimated to different salinity conditions. Error bars represent the standard error of the mean from replicated experiments. Differences between groups were analyzed statistically employing the two-tailed Student’s *t*-test. *P* value < 0.05 was considered statistically significant. * *P* value < 0.05, ** *P* value < 0.01, *** *P* value < 0.001, ns - not significant.

Interestingly, HQ1 worms could acclimate quite well to low salinity condition (0.3% NaCl, referred to as “S3”) which was the standard culture condition for terrestrial *C. elegans*. We observed that, under S3 condition, 2% L1s could develop into adulthood within 3 days, the adulthood percentage was about 92% upon the 7th day, and the earliest egg laying time was around 83 h post L1 stage, which showed no significant differences comparing with S30 group (**Figure 1A, 1B**). However, worms’ lifespan under S3 showed slightly shorter, and L4 worms could live as long as 27 days (**Figure 1C**), with an average lifespan of about 13 days (**Figure 1D**).

Besides, HQ1 worms were able to propagate and acclimate to an even higher salinity environment (5% artificial Sea Salt, referred to as “S50”). Notably, L1 worms could not develop into adult stage within 3 days under S50 condition. The percentage of adulthood upon the 4th and the 5th days also showed significant differences between S50 and S30 groups, while no differences exhibited afterwards (**Figure 1A**). The earliest egg laying time for S50 worms was around 98 h post L1 stage, which showed a severe egg laying delay comparing with both S30 and S3 worms (**Figure 1B**). L4 worms acclimated to S50 condition could live as long as 29 days (**Figure 1C**), with an average lifespan of about 15 days (**Figure 1D**), which was much similar to that of S30 worms.

Taken together, HQ1 worms could acclimate to a wide range of salinity and are one typical euryhaline marine nematodes. How do worms regulate gene expression to acclimate to different salinity environments, remains unknown. Next, we applied newly hatched L1s to investigate the basal differences at both transcriptomic and proteomic levels, respectively.

### Analysis of the Basal Transcriptome for *L. marina* Identifies Genes Involved with Different Salinity Environments

To investigate the basal transcriptomic differences of *L. marina* growing under different salinity environments, we applied newly hatched L1s for RNA-seq analysis. Significant DEGs were identified from different comparison groups (**Figure 2A**), with |log2foldchange| > 1 and DESeq2 padj < 0.05 setting as the differential gene screening thresholds. Details of these DEGs were listed in **Supplementary File 1**.

**FIGURE 2.**
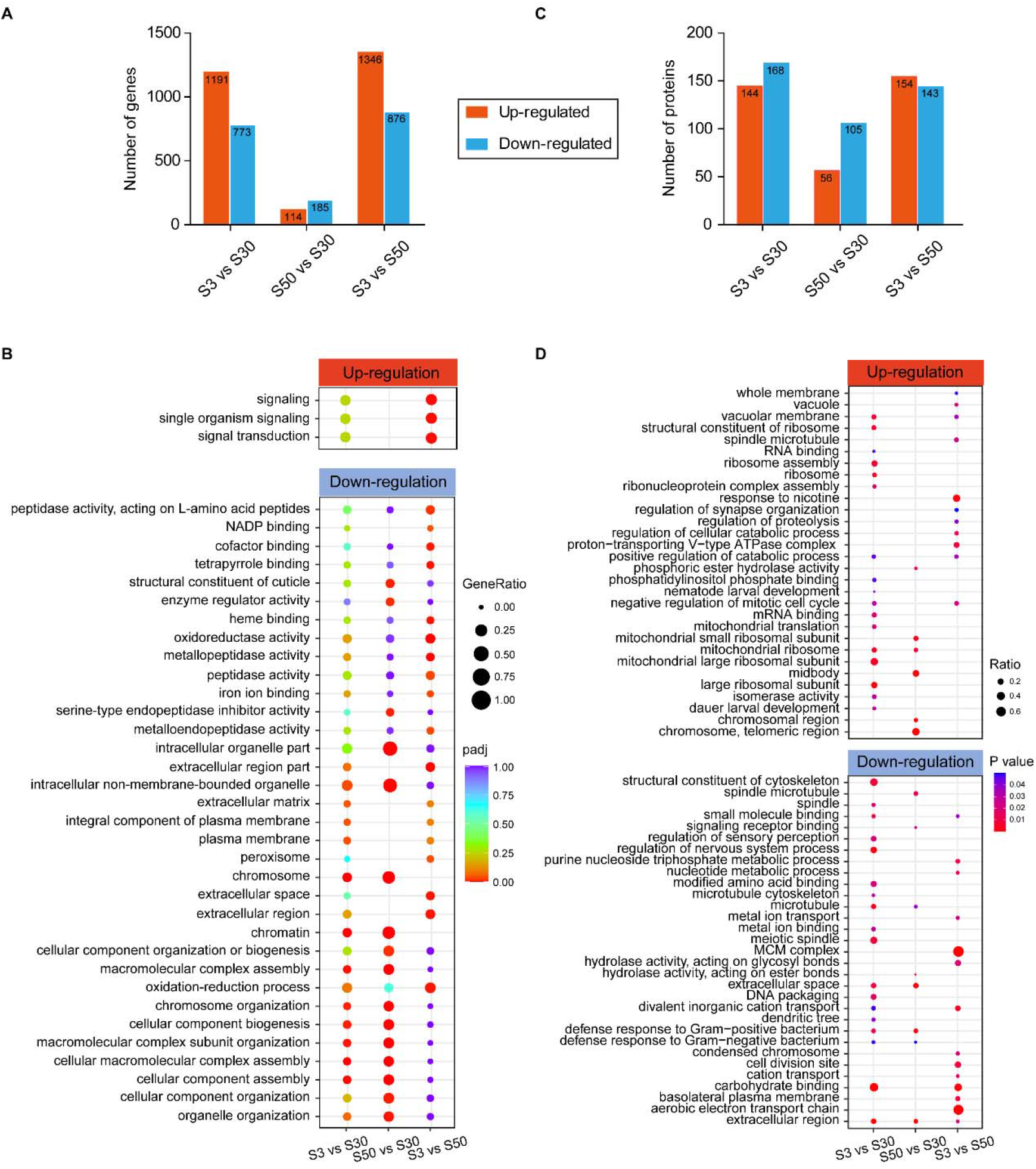
Differentially expressed genes (DEGs) and proteins (DEPs) identified via transcriptomic and proteomic analysis of L1 marine nematodes acclimated to low and high salinity conditions. **(A)** Identified DEGs via RNA-seq analysis for each condition (|log2foldchange| > 1; DESeq2 padj < 0.05 was set as the differential gene screening thresholds). **(B)** Identified DEPs via proteomic analysis for each condition (Ratio of fold change > 1.3 or < 1/1.3; *P* value < 0.05 was set as the differential protein screening thresholds). **(C)** GO enrichment analysis for DEGs. |log2foldchange|>1; DESeq2 padj<0.05 was set as the differential gene screening thresholds. GO enrichment analysis of DEGs were achieved by clusterProfiler R package (v3.4.4), an adjusted *P* value (padj) < 0.05 was considered significantly enriched. The color from red to purple represents the significance of the enrichment. GeneRatio was defined as the ratio of the number of differential genes annotated to the GO term to the total number of differential genes. **(D)** GO enrichment analysis for DEPs. Ratio of fold change > 1.3 or < 1/1.3; *P* value < 0.05 was set as the differential protein screening thresholds. A corrected *P*-value < 0.05 was considered significantly enriched. The color represents the significance of the enrichment. Ratio was defined as the ratio of the number of differential proteins annotated to the GO term to the number of background proteins.

In the low salinity S3 group, a total of 1,191 genes were significantly up-regulated while 773 genes were down-regulated when compared with S30 group (**Figure 2A**). Based on GO enrichment analysis, significant enrichment was only observed within down-regulated DEGs, such genes were mainly annotated to “chromosome” (GO:0005694, padj = 1.88E-06), “cellular macromolecular complex assembly” (GO:0034622, padj = 2.03E-04), “cellular component biogenesis” (GO:0044085, padj = 7.54E-03), “intracellular non-membrane-bounded organelle” (GO:0043232, padj = 3.55E-02), “plasma membrane” (GO:0005886, padj = 3.81E-02) and “extracellular matrix” (GO:0031012, padj = 4.42E-02) (**Figure 2B**). Moreover, “beta-alanine metabolism” (cel00410, padj = 3.83E-02) pathway related genes were significantly enriched among up-regulated DEGs via KEGG pathway enrichment analysis (**Supplementary Figure 4**).

There were relatively less DEGs identified in the S50 versus S30 comparison group compared with S3 versus S30 comparison group (**Figure 2A**). Compared with S30 group, 114 genes were significantly up-regulated in S50 group, of which “purine metabolism” (cel00230, padj = 3.16E-03) pathway related genes were significantly enriched (**Supplementary Figure 4**). In addition, there were 185 DEGs exhibiting significant down-regulation. We found that genes annotated to “chromosome” (GO:0005694, padj = 1.54E-06), “cellular component biogenesis” (GO:0044085, padj = 4.89E-04), “intracellular organelle part” (GO:0044446, padj = 5.72E-04), “intracellular non-membrane-bounded organelle” (GO:0043232, padj = 5.72E-04), “serine-type endopeptidase inhibitor activity” (GO:0004867, padj = 7.96E-03), “structural constituent of cuticle” (GO:0042302, padj = 8.04E-03) and other GO terms were significantly enriched within these down-regulated DEGs (**Figure 2B**).

Additionally, the comparison between S3 group and S50 group was also conducted. There were 1,346 DEGs up-regulated in S3 group (**Figure 2A**), of which “signaling” (GO:0023052, padj = 1.45E-03) related genes were significantly enriched via GO enrichment analysis (**Figure 2B**). It was interestingly noted that a set of small GTPase related genes were included (EVM0002411/*efa-6*, EVM0001914/*exc-5*, EVM0012791/*frm-3*, EVM0004317/*itsn-1*, EVM0014485/*R05G6*.*10*, EVM0004341/*tag-77*, EVM0015120/*tiam-1*, EVM0008542/*unc-73* and EVM0010248/*Y37A1B*.*17*). On the other hand, 876 genes were significantly up-regulated in S50 group (**Figure 2A**), of which “glutathione metabolism” (cel00480, padj = 1.85E-03), “drug metabolism-cytochrome P450” (cel00982,padj = 6.40E-03), “fatty acid degradation” (cel00071, padj = 2.00E-02), “fatty acid metabolism” (cel01212, padj = 2.52E-02) and “metabolism of xenobiotics by cytochrome P450” (cel00980, padj = 2.52E-02) pathway related genes were enriched (**Supplementary Figure 4**). Further, GO enrichment analysis demonstrated that “oxidoreductase activity” genes (GO:0016491, padj = 4.65E-05), “metallopeptidase activity” genes (GO:0008237, padj = 1.74E-04), “extracellular region” genes (GO:0005576, padj = 3.55E-03. Notably, 9 out of the 13 enriched genes were transthyretin-like family genes, TTLs, such as EVM0012475/*ttr-31*, EVM0004658/*ttr-32*, EVM0008749/*ttr-40*, EVM0005297/*ttr-44* and EVM0002004/*ttr-51*.), “heme binding” genes (GO:0020037, padj = 3.15E-03), “cofactor binding” genes (GO:0048037, padj = 5.72E-3), “peptidase activity” genes (GO:0008233, padj = 3.31E-02), “iron ion binding” genes (GO:0005506, padj = 3.47E-02), “peroxisome” genes (GO:0005777, padj = 3.97E-02) and others were significantly enriched within up-regulated DEGs in S50 group (**Figure 2B**).

### Analysis of the Basal Proteome for *L. marina* Identifies Proteins Involved with Different Salinity environments

In parallel, we also applied quantitative proteomic analysis to investigate the basal protein differences among worms growing under different salinity environments. Newly hatched L1s were used, the same developmental staged worms for above transcriptomic analysis. A total of 6,068 proteins were identified (**Supplementary Table 3**) and significant DEPs were selected from different comparison groups (**Figure 2C**), with ratio of fold change > 1.3 or < 1/1.3 and *P* value < 0.05 setting as the differential protein screening thresholds. Details of these DEPs were listed in **Supplementary File 2**.

In the low salinity S3 group, 144 up-regulated proteins and 168 down-regulated proteins were selected comparing with S30 group (**Figure 2C**). Among these up-regulated DEPs, ribosome related proteins were the most significantly enriched ones revealed by both GO enrichment and KEGG pathway enrichment analysis (**Figure 2D** and **Supplementary Figure 5**), others such as “vacuolar membrane” proteins (GO:0005774, *P* value = 6.03E-03), “dauer larval development” proteins (GO:0040024, *P* value = 2.73E-02), “isomerase activity” proteins (GO:0016853, *P* value = 2.83E-02), “negative regulation of mitotic cell cycle” proteins (GO:0045930, *P* value = 3.05E-02), “nematode larval development” proteins (GO:0002119, *P* value = 4.00E-02), “positive regulation of catabolic process” proteins (GO:0009896, *P* value =4.43E-02), “phosphatidylinositol phosphate binding” proteins (GO:1901981, *P* value = 4.47E-02) were also significantly enriched (**Figure 2D**). However, “extracellular region” proteins (GO:0005576, *P* value = 2.29E-06) were the most significantly enriched ones in down-regulated proteins. Besides, cytoskeleton related proteins (including GO:0015630, GO:0005874, **Supplementary Table 4**), defense response related proteins (GO:0050830, *P* value = 1.31E-02; GO:0050829, *P* value = 4.79E-02), “DNA packaging” proteins (GO:0006323, *P* value = 1.71E-02), “modified amino acid binding” proteins (GO:0072341, *P* value = 2.20E-02), “metal ion binding” proteins (GO:0046872, *P* value = 2.69E-02) and “divalent inorganic cation transport” proteins (GO:0072511, *P* value = 4.83E-02) were also significantly enriched (**Figure 2D**).

Compared with S30 group, there were 56 proteins significantly up-regulated in S50 group (**Figure 2C**), of which ribosome related proteins were also significantly enriched via both GO enrichment and KEGG pathway enrichment analysis (**Figure 2D** and **Supplementary Figure 5**). Moreover, “chromosomal region” proteins (GO:0098687, *P* value = 7.33E-04), “midbody” proteins (GO:0030496, *P* value = 7.71E-04, **Supplementary Table 4**) and “phosphoric ester hydrolase activity” proteins (GO:0042578, *P* value = 1.73E-02) were also significantly enriched among these up-regulated DEPs (**Figure 2D**). Additionally, 105 down-regulated DEPs were found in S50 group (**Figure 2C**). Proteins related with “extracellular region” (GO:0005576, *P* value = 2.12E-05) exhibited the most significant enrichment similar to the low salinity S3 group. Besides, microtubule related proteins (GO:0005874, *P* value = 3.87E-02; GO:0005876, *P* value = 1.47E-02, **Supplementary Table 4**), defense response related proteins (GO:0050830, *P* value = 1.21E-03; GO:0050829, *P* value = 4.99E-02) and “hydrolase activity, acting on ester bonds” proteins (GO:0016788, *P* value = 6.87E-03) were also significantly enriched among down-regulated DEPs (**Figure 2D**).

In addition, we also performed the comparison between S3 group and S50 group, and found 154 up-regulated DEPs in S3 group, while 143 up-regulated DEPs in S50 group, respectively (**Figure 2C**). Based on KEGG pathway enrichment analysis, “lysosome” pathway related proteins (map04142, *P* value = 7.09E-06) and “DNA replication” pathway related proteins (map03030, *P* value =1.75E-05) were the most significantly enriched ones in S3 and S50 groups, respectively (**Supplementary Figure 5**). Besides, based on GO enrichment analysis, “response to nicotine” proteins (GO:0035094, *P* value = 2.46E-03), “proton-transporting V-type ATPase complex” proteins (GO:0033176, *P* value = 8.38E-03, including EVM0007934/VHA-5 and EVM0016316/VHA-6), “whole membrane” proteins (GO:0098805, *P* value = 4.83E-02), “regulation of cellular catabolic process” proteins (GO:0031329, *P* value = 1.11E-02), “spindle microtubule” proteins (GO:0005876, *P* value = 2.20E-02, **Supplementary Table 4**), “negative regulation of mitotic cell cycle” proteins (GO:0045930, *P* value = 2.28E-02), “regulation of proteolysis” proteins (GO:0030162, *P* value = 3.92E-02), “regulation of synapse organization” proteins (GO:0050807, *P* value = 4.86E-02) exhibited significant enrichment in S3 group (**Figure 2D**). In contrast, “MCM complex” proteins (GO:0042555, *P* value = 2.11E-05), “condensed chromosome” proteins (GO:0000793, *P* value = 2.36E-02), “extracellular region” proteins (GO:0005576, *P* value = 2.21E-02), “cation transport” proteins (GO:0006812, *P* value = 1.49E-02), “nucleotide metabolic process” proteins (GO:0009117, *P* value = 1.18E-02), “cell division site” proteins (GO:0032153, *P* value = 1.05E-02), “hydrolase activity, acting on glycosyl bonds” proteins (GO:0016798, *P* value = 1.96E-02) and others were significantly enriched in S50 group (**Figure 2D**).

### Identification of Genes Expressed Consistently at both mRNA and Protein Levels in Different Salinity Environments

Interestingly, not only salinity differences between S50 and S30 conditions was smaller than that of S3 versus S30 conditions, but we also noticed a similar tendency, based on the number of DEGs and DEPs identified within different comparisons, that the differences between S50 and S30 were smaller when compared with that from the S3 versus S30 comparison pair, especially for DEGs. Hereby, we refer to these worms which were acclimated to S3 environment as “low salinity group”, while worms growing under S30 and S50 environments as “high salinity groups”. Further, in order to identify crucial genes related with euryhalinity, we tried to screen genes that were expressed consistently at both mRNA and protein levels.

As shown in Figure 3A, 1,638 genes were demonstrated up-regulation specifically in low salinity group (S3) compared with high salinity groups (S30 and S50), with screening thresholds set to fold change > 1.0, padj < 0.05 applied to transcriptomic data. Similarly, with screening thresholds set to fold change > 1.0, *P* value < 0.05 applied to proteomic data, a total of 354 proteins were specifically selected with up-regulation in low salinity group. Furthermore, we combined the results from transcriptomic and proteomic analysis and found that there were 78 genes exhibiting consistent expression at both mRNA and protein levels. Interestingly, we also found that 29 out of the above 78 genes, including the trehalose biosynthesis gene EVM0016452/*tps-2*, the transthyretin-like family genes EVM0003972/*ttr-15* and EVM0004638/*ttr-30*, the ion transport genes EVM0007934/*vha-5*, EVM0000094/*twk-33* and EVM0008374/*mca-1*, demonstrated specific induction upon hyposaline stress in *L. marina* based on our previous data (**Supplementary Table 5**). Thus, these 78 genes were considered as low salinity specific genes, and their detailed information was summarized in Table 1.

**FIGURE 3.**
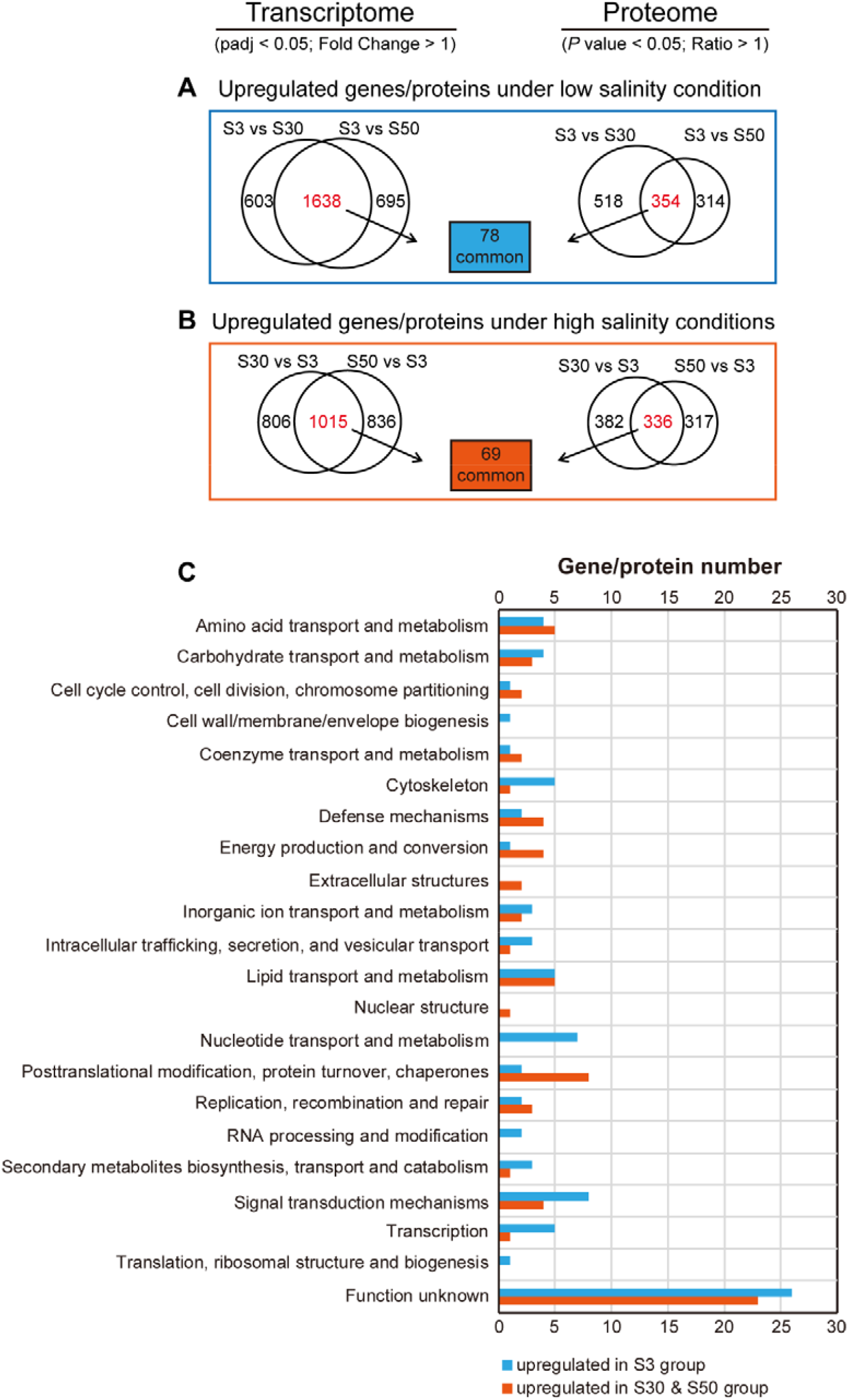
Identification of genes expressed consistently at both mRNA and protein levels under low and high salinity conditions. (**A**) Identification of 78 genes with upregulated expression at both mRNA and protein levels under low salinity condition (S3 group) compared with high salinity conditions (S30 and S50 groups). DESeq2 padj < 0.05; Fold Change > 1 was set as the differential gene screening threshold. *P* value < 0.05; Ratio > 1 was set as the differential protein screening threshold. (**B**) Identification of 69 genes with upregulated expression at both mRNA and protein levels under high salinity conditions (S30 and S50 groups) compared with low salinity condition (S3 group). DESeq2 padj < 0.05; Fold Change > 1 was set as the differential gene screening threshold. *P* value < 0.05; Ratio > 1 was set as the differential protein screening threshold. (**C**) Classification of differential genes expressed under low and high salinity conditions based on COG annotation.

**TABLE 1.**
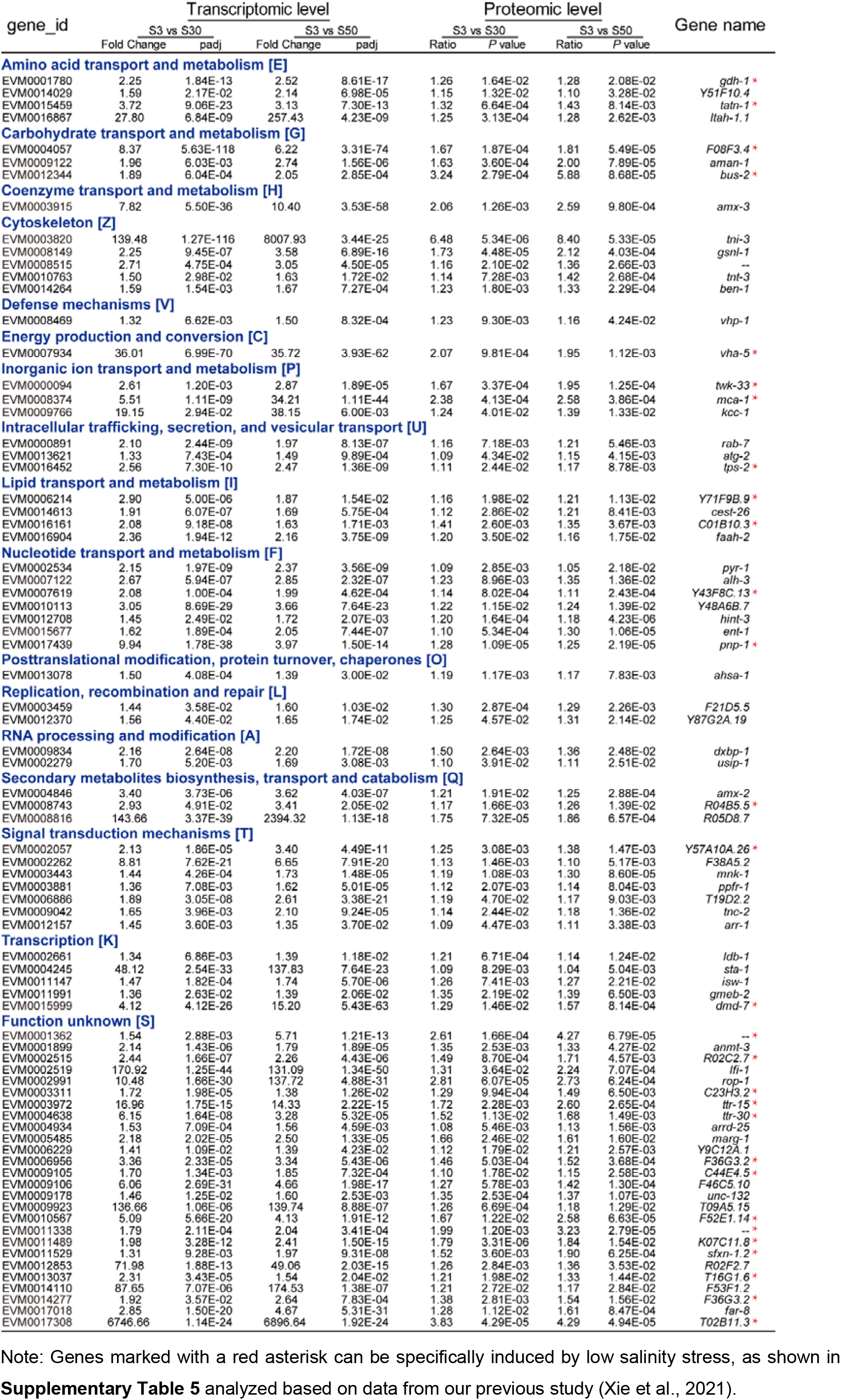
Detailed information for 78 Common DEGs and DEPs up-regulated specifically under low salinity.

To identify high salinity specific genes, similar analysis was performed as described above. Briefly, 1,015 genes and 336 proteins were specifically up-regulated in high salinity groups (S30 and S50), respectively (**Figure 3B**). A total of 69 genes were further selected as high salinity specific genes, with detailed information summarized in Table 2. Additionally, there were 11 genes among these 69 genes, including the glycerol biosynthesis gene EVM0001663/*gpdh-1* and the cuticle related collagen gene EVM0013254/*col-160*, can be specifically induced by hypersaline stress (**Supplementary Table 6**).

**TABLE 2.**
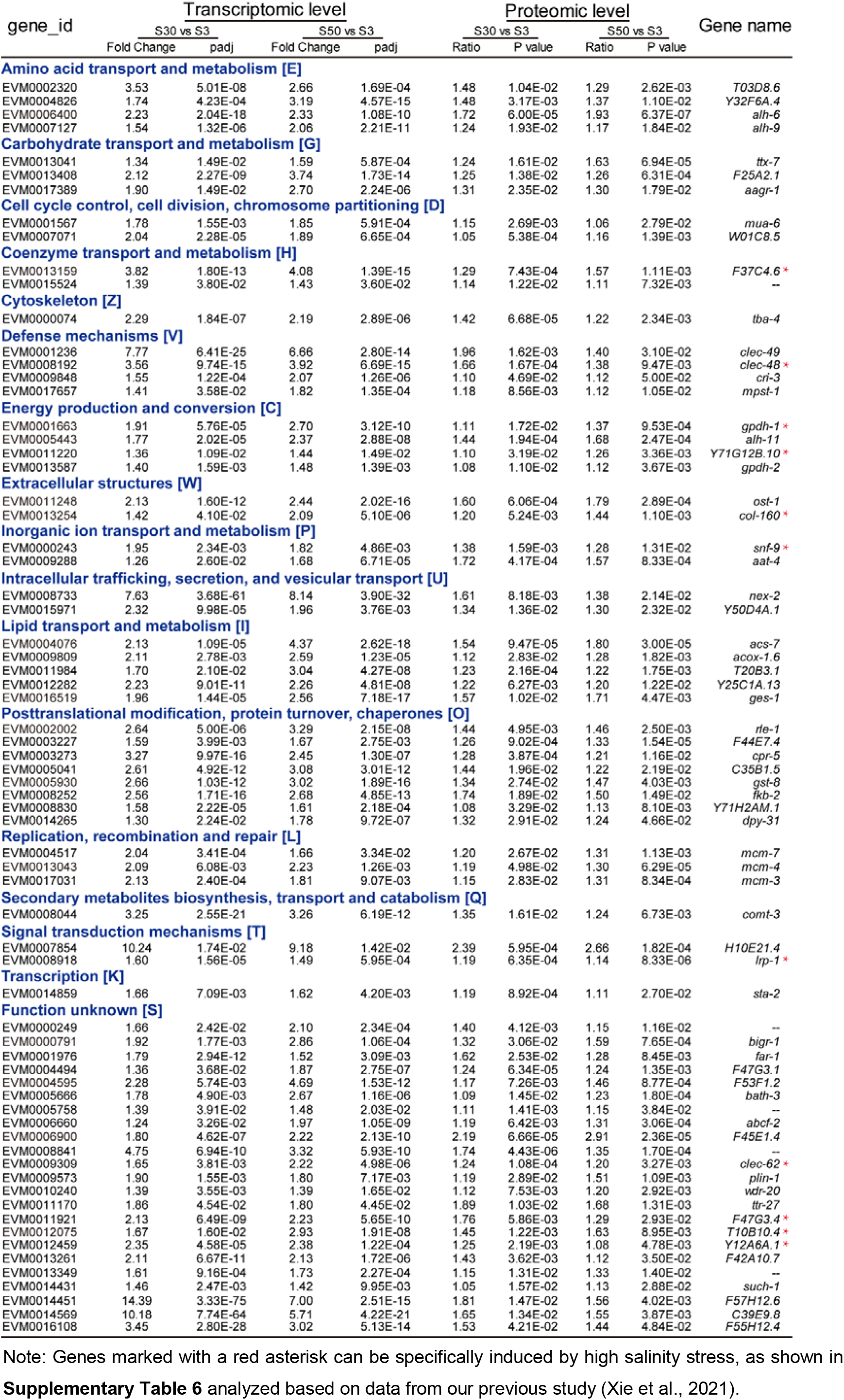
Detailed information of 69 Common DEGs and DEPs up-regulated specifically under high salinity environments.

Further, the differential genes expressed specifically under low and high salinity conditions were classified based on their annotation information (**Figure 3C**). We found that, under low salinity condition, the top six genes/proteins categories with annotated function were grouped to “signal transduction mechanisms” (EVM0003443/*mnk-1*, EVM0003881/*ppfr-1*, EVM0009042/*tnc-2* and EVM0012157/*arr-1*), “nucleotide transport and metabolism” (EVM0002534/*pyr-1*, EVM0007122/*alh-3*, EVM0015677/*ent-1*, EVM0017439/*pnp-1* and EVM0012708/*hint-3*), “cytoskeleton” (EVM0003820/*tni-3*, EVM0008149/*gsnl-1*, EVM0010763/*tnt-3* and EVM0014264/*ben-1*), “lipid transport and metabolism” (EVM0014613/*cest-26*, EVM0016904/*faah-2* and EVM0016867/*ltah-1*.*1*), as well as “transcription” (EVM0002661/*ldb-1*, EVM0004245/*sta-1*, EVM0011991/*gmeb-2*, EVM0015999/*dmd-7* and EVM0011147/*isw-1*) (**Table 1**). However, “posttranslational modification, protein turnover, chaperones” (EVM0002002/*rle-1*, EVM0003273/*cpr-5*, EVM0005930/*gst-8*, EVM0008252/*fkb-2* and EVM0014265/*dpy-31*), “amino acid transport and metabolism” (EVM0006400/*alh-6* and EVM0007127/*alh-9*), “lipid transport and metabolism” (EVM0004076/*acs-7*, EVM0009809/*acox-1*.*6* and EVM0016519/*ges-1*), “defense mechanisms” (EVM0009848/*cri-3*, EVM0017657/*mpst-1*, EVM0008192/*clec-48* and EVM0001236/*clec-49*), “energy production and conversion” (EVM0001663/*gpdh-1*, EVM0013587/*gpdh-2* and EVM0005443/*alh-11*), as well as “signal transduction mechanisms” (EVM0008918/*lrp-1*, etc.) related genes/proteins were the top six categories under high salinity conditions (**Table 2**).

### Identification of Genes and Their Corresponding Proteins Whose Abundance was Proportional to Environmental Salinity

In the present study, we found that DEGs enriched markedly based on RNA-seq analysis, actually hardly exhibited enrichment at the protein level (**Figure 2B** and **2D**), and it is the fact that only limited genes expressed consistently at both mRNA and protein levels (**Figure 3A** and **3B**). Next, we focused on those genes whose abundance was directly or inversely proportional to environmental salinity at both mRNA and protein levels, that were probably key regulators or effectors associated with its effective osmoregulation of euryhaline *L. marina*.

We therefore combined transcriptomic and proteomic profiles and further found that 66 genes and their corresponding proteins demonstrated environmental salinity dependent patterns in expression (**Supplementary File 3**). Specifically, 38 genes were down-regulated when salinity increased, while 28 genes were up-regulated when salinity increased (**Figure 4** and **Supplementary File 3**).

**FIGURE 4.**
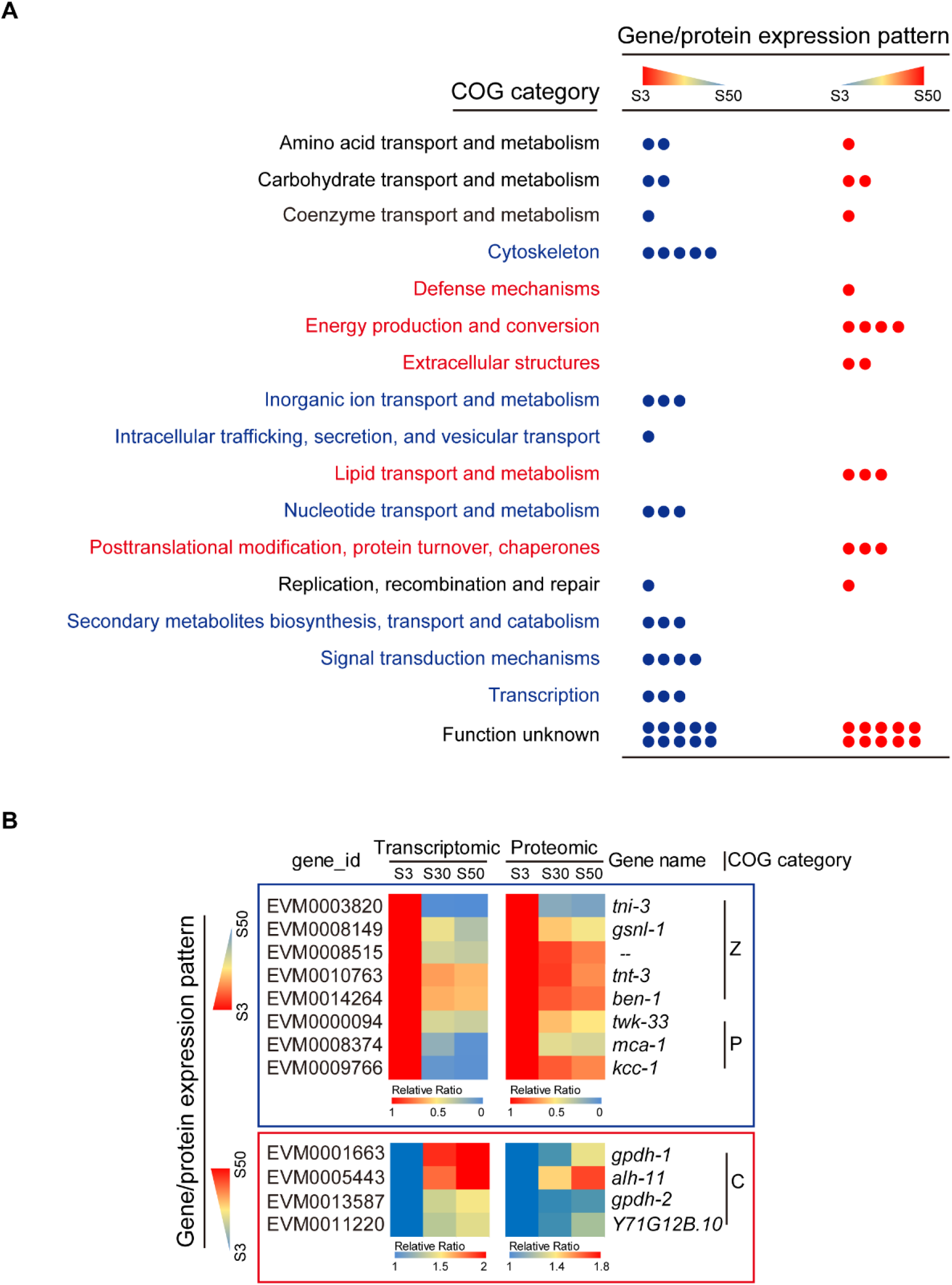
Identification of genes and their corresponding proteins whose abundance was directly or inversely proportional to environmental salinity. (**A**) Differential genes which were expressed in a salinity dependent pattern were classified based on COG annotation. Each dot represents an individual gene. Blue: Genes/proteins that down-regulated in expression with increasing salinity. Red: Genes/proteins that up-regulated in expression with increasing salinity. Detailed information can be found in **Supplementary file 3**. (**B**) Expression profiles of representative genes/proteins with opposite pattern.

We classified the candidates by their function annotation information and found that, interestingly, “cytoskeleton” related genes/proteins (EVM0003820/*tni-3*, EVM0008149/*gsnl-1*, EVM0010763/*tnt-3*, EVM0014264/*ben-1* and EVM0008515), “inorganic ion transport and metabolism” related genes/proteins (EVM0000094/*twk-33*, EVM0008374/*mca-1* and EVM0009766/*kcc-1*), “signal transduction mechanisms” related genes/proteins (EVM0003443/*mnk-1*, EVM0003881/*ppfr-1* and EVM0009042/*tnc-2*), as well as “transcription” related genes/proteins (EVM0011147/*isw-1*, EVM0011991/*gmeb-2* and EVM0015999/*dmd-7*) and others (“intracellular trafficking, secretion, and vesicular transport”, “nucleotide transport and metabolism” and “secondary metabolites biosynthesis, transport and catabolism”) were notably grouped showing up-regulation with the decreasing salinity (**Figure 4A** and **4B**). On the contrary, “energy production and conversion” related genes/proteins (EVM0001663/*gpdh-1*, EVM0013587/*gpdh-2*, EVM0005443/*alh-11* and EVM0011220/*Y71G12B*.*10*), “lipid transport and metabolism” related genes/proteins (EVM0004076/*acs-7*, EVM0009809/*acox-1*.*6* and EVM0016519/*ges-1*), as well as “posttranslational modification, protein turnover, chaperones” related genes/proteins (EVM0002002/*rle-1*, EVM0005930/*gst-8* and EVM0008830/*Y71H2AM*.*1*), and others (“extracellular structures” and “defense mechanisms”) were grouped exhibiting up-regulation with the increasing salinity (**Figure 4A** and **4B**).

Together, the above genes could be key regulators or effectors in *L. marina* involved in its acclimation process to different salinity environments, which were worthy of further in-depth functional study in the future.

## Discussion

Environmental salinity is particularly critical for aquatic animals. Effective osmotic regulation not only directly affects their survival, but also shapes their behavior and distribution. Studies in euryhaline fish have provided important insights into the mechanisms of osmotic response and adaptation for multicellular organisms. There are various euryhaline fish in nature, such as eels spending most of their lives in freshwater until they return to their spawning grounds in the sea, whereas salmons migrating from ocean through the natal river for spawning, other fish, for example, Atlantic killifish (*Fundulus heteroclitus*), Arabian pupfish (*Aphanius dispar*) and threespine sticklebacks (*Gasterosteus aculeatus*) are widely distributed in freshwater as well as brackish and marine areas (Whitehead et al., 2011; Jones et al., 2012; Bonzi et al., 2021). During freshwater to seawater transition or vice versa, euryhaline fish although cope with external salinity changes in a species-specific way, evolutionary conserved strategies do exist among them. The gill, intestine and kidney are the major osmoregulatory tissues in fish, successful acclimation in both freshwater and seawater environments depends on proper physiological, metabolic and structural adjustments in these tissues, which also involves their neuroendocrine regulation (Kültz, 2012; 2015). Generally, fish increase intestinal water absorption and salt secretion in gill under hypersaline conditions, by contrast, the fish gill reabsorbs salt and the kidney produces diluted urine under hyposaline conditions. Euryhaline fish have the ability to perceive salinity changes through multiple osmo-sensors, including transmembrane proteins and cytoskeletal proteins, which results in an early osmotic response and regulation processes, the following regulatory expression and activities of diverse genes and corresponding proteins involved in water and ions transport, macromolecular damage control, cell cycle process, apoptosis, accumulation and transport of organic osmolytes, reallocation of energy and other physiological processes together contribute to cellular stress response, cell volume regulation as well as tissue remodeling (Copeland, 1950; Tseng and Hwang, 2008; Whitehead et al., 2011; Kültz, 2015; Evans and Kültz, 2020). In addition to the final physiological acclimation to external salinity, there are also phenotypic differences in behavior and body morphology or size between marine populations and freshwater populations as reported in *Fundulus* and threespine sticklebacks (Kültz et al., 2015; Styga et al., 2019). Similar regulatory mechanisms have also been reported in crustaceans, like crabs and shrimps (Thabet et al., 2017; Rahi et al., 2019; Niu et al., 2020). However, the precise mechanisms underlying osmotic sensation, signal transduction and adaptation remain elusive. In this study, we applied a newly emerging model system, marine nematode *L. marina*, and analyzed the basal transcriptomic and proteomic differences among worms acclimated to different salinities, aiming to provide insight into invertebrate euryhalinity.

Based on our studies, marine nematode *L. marina* can tolerate and acclimate to a wide range of salinity and is a typical euryhaline marine invertebrate. In an effort to identify salinity responding genes in *L. marina*, we previously applied newly hatched L1 larvae and stressed them by either low or high salinity conditions. We observed that worms were paralyzed immediately in parallel with obvious body volume alterations upon both low and high salinity stresses, and these stressed worms both exhibited developmental defects afterwards (Xie et al., 2021). In the present study, worms could move, grow and propagate normally after acclimation to either hyposaline or hypersaline environments for a long time, except that a relatively shortened lifespan for hyposaline acclimated worms and a relatively delayed development for hypersaline acclimated worms. Studies in fish and crustaceans had revealed that osmoregulatory epithelia, in particular the gill epithelium, are remodeled during the transition between freshwater and seawater environments (Copeland, 1950; Thabet et al., 2017; Evans and Kültz, 2020). However, we are not sure whether there are any differences regarding specific tissue structures or body size between those worms acclimated to different salinity conditions, which await elucidation afterwards. Nevertheless, candidate genes involved in osmotic regulation and euryhaline acclimation were identified from our studies at the organismal level in *L. marina*.

### Accumulation of Organic Osmolytes is Required for both Early Salinity Stress Response and Long-term Salinity Acclimation in *L. marina*

It is known that the accumulation of organic osmolytes is a ubiquitous mechanism in cellular osmoregulation (Yancey, 2005; Burg and Ferraris, 2008). There are a number of organic osmolytes such as glycerol, trehalose, inositol, betaine and taurine, allow cells to against effects of hyperosmolarity and to adapt to hyperosmotic conditions. For example, free amino acids and methylamines are mainly utilized by most marine invertebrates (Thabet et al., 2017; Niu et al., 2020), whereas glycerol is the most important osmolyte for the terrestrial nematode *C. elegans* (Choe, 2013; Urso and Lamitina, 2021). It is well documented that an increased level of glycerol is essential for *C. elegans’* survival in hypertonic environments mediated by upregulation of the glycerol biosynthetic enzyme gene *gpdh-1* (Lamitina et al., 2006; Urso and Lamitina, 2021). Burton et al. reported that if *C. elegans* was exposed to mild high salinity stress, its progeny could be protected from strong osmotic stress via increasing the expression of *gpdh-2*, and this glycerol synthesis gene is essentially required for the transgenerational osmotic protection (Burton et al., 2017). In line with its terrestrial relative *C. elegans*, we previously demonstrated that *gpdh-1* were significantly induced by high salinity stress in *L. marina*, suggesting that glycerol as an essential osmolyte in both nematodes’ response to hyperosmotic stress (Xie et al., 2021). In the current study, two glycerol synthesis genes, EVM0001663/*gpdh-1* and EVM0013587/*gpdh-2*, both showed significant up-regulation in high salinity groups at both mRNA and protein levels (**Table 2** and **Figure 4B**), whose expression was directly proportional to the level of salinity. These data suggest that *L. marina* may use glycerol as an osmolyte in both response and acclimation to high salinity environments. Given *gpdh-2* was not notably induced by salinity stress, but was involved in hypersaline acclimation, our results indicate that *gpdh-2* might play major roles in transgenerational osmotic protection inheritance and hyperosmotic acclimation in the marine nematode *L. marina*. Additionally, we also noticed a STAT transcription factor-like gene EVM0014859/*sta-2* expressed in consistent with EVM0001663/*gpdh-1* under high salinity conditions (**Table 2**). As reported by Dodd et al. (Dodd et al., 2018), RNAi of *sta-2* decreased *gpdh-1* reporter induction in *dpy-7* mutants, indicating *sta-2* plays a positive role in glycerol accumulation in *C. elegans*. Here, based on the consistent expression pattern between EVM0001663/*gpdh-1* and EVM0014859/*sta-2* in this context, we thus proposed a conserved transcriptional regulation of *sta-2* on *gpdh-1* expression in *L. marina*, which perhaps be involved in adaptive regulation under hypersaline conditions. Moreover, we previously reported that trehalose biosynthesis gene EVM0007411/*tps-2* was induced by both low and high salinity stresses in *L. marina* (Xie et al., 2021). Here, we found that two annotated *L. marina tps-2* genes, EVM0007411 and EVM0016452, both significantly elevated their transcription under low salinity. Their overall protein levels were also increased, especially for EVM0007411, which showed notably up-regulation in S3 worms when compared with S50 worms (**Supplementary Table 7**).

Taken together, we concluded that both trehalose and glycerol are required not only for short-term salinity stress response but also for long-term salinity acclimation. On the other hand, trehalose potentially contributes to hyposaline acclimation while glycerol most probably accompanies hypersaline acclimation in *L. marina*.

### Cellular Stress Response in Osmotic Regulation of *L. marina*

Several basic aspects of the osmotic induced cellular stress response were well documented, including regulation on cell cycle and reallocation of metabolic energy (Evans and Kültz, 2020). Here, diverse genes related with “cell cycle control, cell division, chromosome partitioning” and “replication, recombination and repair”, were identified in both low and high salinity acclimated groups, as summarized in Figure 3C. Similarly, metabolism related genes associated with “amino acid transport and metabolism”, “carbohydrate transport and metabolism”, “coenzyme transport and metabolism”, “lipid transport and metabolism”, “nucleotide transport and metabolism” and “energy production and conversion”, also exhibited differential expression patterns in worms growing under hyposaline and/or hypersaline acclimated conditions (**Figure 3C**). Moreover, we interestingly found that many of these genes, especially for metabolism related ones, could also be induced by early short-term salinity stresses, for instance, EVM0001780/*gdh-1*, EVM0015459/*tatn-1*, EVM0012344/*bus-2*, EVM0004057/*F08F3*.*4*, EVM0016161/*C01B10*.*3*, EVM0006214/*Y71F9B*.*9*, EVM0017439/*pnp-1* and EVM0007619/*Y43F8C*.*13* were responsive to hyposaline stress (**Table 1** and **Supplementary Table 5**), while EVM0001663/*gpdh-1*, EVM0011220/*Y71G12B*.*10* and EVM0013159/*F37C4*.*6* were responsive to hypersaline stress (**Table 2** and **Supplementary Table 6**), suggesting the involvement of these genes in both osmotic stress response and acclimation processes in euryhaline *L. marina*.

Another aspect of the osmotic induced cellular stress response is programmed cell death (Evans and Kültz, 2020). Previously, we reported a group of TTL genes presumably involved in apoptosis and the damage control regulation in response to both low and high salinity stresses in *L. marina* (Xie et al., 2021). Here, we found that diverse TTL genes were differentially expressed in acclimation to different salinity environments. Totally 31 out of about 47 annotated TTL genes in *L. marina*’s genome exhibited expression differences with significance in at least one comparison group at either mRNA or protein level or both (**Supplementary Table 8**). For instance, 10 TTL genes upregulated under low salinity condition, with 4 only increased abundance at mRNA level (EVM0010840/*ttr-15*, EVM0004159/*ttr-59*, EVM0009470/*ttr-53* and EVM0016217/*ttr-54*) and 2 only increased abundance at protein level (EVM0003838/*ttr-46*, EVM0005032/*ttr-7*), another 4 increased abundance at both levels (EVM0016470/*ttr-59*, EVM0008626/*ttr-46*, EVM0003972/*ttr-15* and EVM0004638/*ttr-30*). By contrast, 17 TTL genes upregulated under high salinity condition(s), with 9 only increased abundance at mRNA level (EVM0004800/*ttr-16*, EVM0003534/*ttr-15*, EVM0016920/*ttr-18*, EVM0012475/*ttr-31*, EVM0014605/*ttr-8*, EVM0010105*/ttr-44*, EVM0010790/*ttr-33*, EVM0006931/*ttr-49* and EVM0008749/*ttr-40*) and 4 only increased abundance at protein level (EVM0003282/*ttr-41*, EVM0001487/*ttr-27*, EVM0008675/*ttr-5* and EVM0015122/*ttr-36*), another 4 increased abundance at both levels (EVM0002004/*ttr-51*, EVM0005297/*ttr-44*, EVM0007550/*ttr-30* and EVM0011170/*ttr-27*). In addition, there were 4 other genes demonstrated opposing changes at mRNA level and protein level (Under low salinity, EVM0004593/*ttr-27* increased mRNA level but decreased protein abundance, whereas EVM0005705/*ttr-5*, EVM0017357/*ttr-44* and EVM0004658/*ttr-32* decreased mRNA levels but increased protein abundance.). Together, these distinctively differential regulation of diverse TTL genes in either hyposaline or hypersaline acclimated nematodes, suggesting that certain TTL genes might play essential roles in hyposaline acclimation, while others play major roles in hypersaline acclimation. Moreover, 16 out of the above 31 salinity acclimation related TTL genes have been identified in our previous study, which could also be induced by both hypo- and hyper-osmotic stresses, such as EVM0004638/*ttr-30*, EVM0005297/*ttr-44*, EVM0008626/*ttr-46* and EVM0016470/*ttr-59* (**Supplementary Table 8**). As one of the largest nematode-specific protein families, most TTLs were predicted to be secreted (Parkinson et al., 2004;Wang et al., 2010). Some TTL genes in *C. elegans* were responsive to diverse environmental challenges including oxidative stress, pathogen exposure and osmotic stress (Oliveira et al., 2009; Engelmann et al., 2011; Schmeisser et al., 2013; Treitz et al., 2015; Dodd et al., 2018). According to studies on *ttr-52*, which was reported as a bridging factor involved in cell corps engulfment, apoptosis and axon repair (Wang et al., 2010; Mapes et al., 2012; Neumann et al., 2015), and another TTL gene *ttr-33*, which was reported as a potential secreted sensor or scavenger of oxidative stress involved in neuroprotective mechanism (Offenburger et al., 2018). We thus speculated that TTL genes, presumably function in apoptosis and damage control machinery involved in the cellular stress response to cope with salinity stresses, and play important roles in both salinity response and acclimation processes in euryhaline *L. marina*. How each TTL gene functions underlying salinity acclimation deserves to be further explored.

### Cell Volume Regulation in Osmotic Regulation of *L. marina*

Cell volume is known to be affected by osmotic exposure. Profound changes, such as cell shrinkage and swelling, can be observed upon acute hyperosmotic and hypoosmotic stresses, respectively. Volume-dependent regulation will be evolved to restore near-normal cell volume to maintain homeostasis after volume perturbation (Hoffmann et al., 2009). The cytoskeleton was implicated as a potential volume sensor and a mediator of volume-dependent regulation of various ion transporters and channels (Henson, 1999; Pedersen et al., 2001; Hoffmann et al., 2009). It has been well documented that cytoskeletal remodeling, especially actin-related, is involved in volume changes upon osmotic stress (Di Ciano et al., 2002; Di Ciano-Oliveira et al., 2006). Previously, we reported that several tubulin genes show significantly opposing transcriptional changes between low and high salinity stressed worms, indicating their potential roles in salinity response in *L. marina* (Xie et al., 2021). In the present study, although some tubulin related proteins were enriched in different salinity comparison groups based on the proteomic data, they actually hardly expressed consistently at the mRNA level (**Figure 2D** and **Supplementary Table 4**). However, five cytoskeleton related genes, EVM0003820/*tni-3*, EVM0008149/*gsnl-1*, EVM0010763/*tnt-3*, EVM0014264/*ben-1* and EVM0008515, exhibited remarkable elevation at both mRNA and protein levels in low salinity acclimated environment (**Table 1**). Of note, expression of all these five genes was inversely proportional to the environmental salinity, increasing their abundance as salinity decreases (**Figure 4B**), suggesting their important roles in hyposaline acclimation in *L. marina*. Three of these are actin regulatory genes, for example, *tni-3* and *tnt-3* are orthologs of human troponin I and troponin T respectively (Meissner et al., 2009; Takashima et al., 2012), *gsnl-1* is an ortholog of human gelsolin family gene (Klaavuniemi et al., 2008; Liu and Ono, 2013). Both troponin and gelsolin are involved in regulation of actin dynamics in a calcium-dependent manner, playing important roles in various actin-related processes, including cell structure, cell growth, cell motility, intracellular transport and muscle contraction (Gordon et al., 2000; Ono, 2007; Nag et al., 2013). Although this is the first time exhibiting a potential role for troponin and gelsolin related proteins in osmoregulation in *L. marina* or even in marine invertebrates, there were already numerous evidence upon their involvement in various human diseases from inflammation to cancer (Li et al., 2012; Welsh et al., 2019; Imazio et al., 2020). Hereby, we are curious about whether cellular or muscular defects or remodeling accompany worm’s acclimation to either low or high salinity environments. Detailed cellular, molecular and genetic studies on how these actin regulatory genes function in euryhaline acclimation in *L. marina* deserves to be further explored.

Interestingly, corresponding with expression profiles of these actin regulatory genes, one plasma membrane calcium pump gene EVM0008374/*mca-1* (Carafoli, 1987; Kraev et al., 1999) also exhibited elevation in both mRNA and protein abundances when salinity decreased (**Figure 4B**). As an important component for the maintenance of calcium homeostasis in cells, *mca-1* probably has a role in actin cytoskeleton regulation in *L. marina*. Besides, Rho family small GTPases have been suspected to participate in the osmotically induced remodeling of the cytoskeleton (Pedersen et al., 2001; Di Ciano-Oliveira et al., 2006). Consistent with these findings, we also found a set of small GTPase related genes via GO enrichment analysis (GO:0023052, “signaling”, padj = 1.45E-03; GO:0035023, “regulation of Rho protein signal transduction”, *P* value = 0.00317; GO:0007264, “small GTPase mediated signal transduction”, *P* value = 0.016904), which only increased their transcriptional levels under low salinity condition (**Supplementary Table 9**), yet might indicating their potential involvement in cytoskeleton regulation associating with hyposaline acclimation in *L. marina*.

In marine invertebrates and fish, ion transporters and channels play important roles in osmoregulation (Kültz, 2015; Thabet et al., 2017; Niu et al., 2020; Vij et al., 2020; Zhang et al., 2020). One example is the upregulation of V-type H^+^-transporting ATPase genes in response to low salinity stress, which have been documented not only in the crustacean like shrimp *Litopenaeus vannamei* (Wang et al., 2012) and the mud crab *Scylla paramamosain* (Niu et al., 2020), but also in marine nematode *L. marina* (Xie et al., 2021), indicating their conserved function among marine invertebrates. Besides, Bonzi et al. (Bonzi et al., 2021) reported that ion transport genes showed significant expression differences in gill tissues between two natural populations of Arabian pupfish (*Aphanius dispar*), which inhabiting nearly-freshwater and seawater areas, respectively. In our present study, in addition to the V-type H^+^-transporting ATPase gene EVM0007934/*vha-5* and a calcium pump gene EVM0008374/*mca-1* discussed above, expression of a potassium channel gene EVM0000094/*twk-33* (Buckingham et al., 2005) and a potassium/chloride cotransporter gene EVM0009766/*kcc-1* (Holtzman et al., 1998), was also found inversely proportional to the level of salinity, elevating both mRNA and protein abundances when salinity was decreasing (**Figure 4B**). The increased abundance of potassium channel and potassium/chloride cotransporter genes was consistent with their general requirement mediating K^+^ and Cl^-^ efflux to defense against cell swelling under hypoosmotic (Strange, 2004; Hoffmann et al., 2009). Specifically, it has long been documented that the potassium/chloride cotransporters regulate a broad range of physiological processes, including cell volume regulation and ion homeostasis in many vertebrate cell types, and are among the most medically relevant ion transporters in human (Lauf and Theg, 1980; Adragna et al., 2004; Kahle et al., 2015; Meor Azlan and Zhang, 2020; Zimanyi et al., 2020). Moreover, EVM0007934/*vha-5*, EVM0008374/*mca-1* and EVM0000094/*twk-33*, were also demonstrated specific induction upon hyposaline stress in our previous report (Xie et al., 2021) (**Table 1** and **Supplementary Table 5**), we therefore proposed that these above-mentioned ion transport related genes play crucial roles in both osmotic stress response and long-term acclimation processes specifically under low salinity condition in *L. marina*.

Additionally, we previously found strong positive selections on six genes including EVM0011198/*col-109*, EVM0009368/*lec-1*, EVM0011012/*T13H5*.*6*, EVM0014994/*osm-12*, EVM0011797/*gem-1*, and EVM0008924/*imp-2* in *L. marina* genome (Xie et al., 2020), which highlighted potential roles related with this nematode’s adaptation to a marine habitat. In a more recent study, we did find that four of these putative salinity-related positive selected genes (PSGs) were significantly regulated in response to salinity stresses (Xie et al., 2021). In the current study, EVM0011198/*col-109* and EVM0009368/*lec-1* were demonstrated alteration in expression either at mRNA or protein levels with significance among different salinity acclimated groups (**Supplementary Table 10**). Further genetic and cellular analysis of these PSGs in *L. marina* should establish their functions in osmoregulation.

## Conclusion

In conclusion, we have described for the first time the genome-wide transcriptional and proteomic analysis of the marine nematode *L. marina* acclimated to either low salinity or high salinity conditions. We found that various cellular stress response genes may function as the conserved regulators in both short-term salinity stress response and long-term acclimation processes. Additionally, we identified diverse genes involved in cell volume regulation which might contribute to hyposaline acclimation, and we also demonstrated that genes related to glycerol biosynthesis might accompany hypersaline acclimation in *L. marina*. Moreover, our results indicated that the glycerol biosynthesis gene *gpdh-2* probably plays a major role in transgenerational inheritance of osmotic stress protection in worms. Thus, our data might lay the foundation to identify the key gene(s) for further in-depth exploration on environmental adaptation mechanisms in euryhaline organisms, especially in the context of global climate change and the corresponding marine salinity stresses.

### Article types

Original Research Article.

## Author Contributions

YX and LZ conceived and designed the experiments. YX carried out the experiments, analyzed the data, and wrote the manuscript. LZ edited the manuscript and supervised the project. Both authors read and approved the final manuscript.

## Acknowledgments

We are grateful to all members of the LZ laboratory for their helpful discussions.

## Data Availability Statement

The RNA-seq raw data were submitted to the NCBI under the accession code PRJNA778902. All raw proteomics data are available via ProteomeXchange under identifier PXD029671.

